# Brain Morphometry and Diminished Physical Growth in Bangladeshi Children Growing up in Extreme Poverty: a Longitudinal Study

**DOI:** 10.1101/2021.02.24.432797

**Authors:** Ted K. Turesky, Talat Shama, Shahria Hafiz Kakon, Rashidul Haque, Nazrul Islam, Amala Someshwar, William A. Petri, Charles A. Nelson, Nadine Gaab

## Abstract

Diminished physical growth is a common marker of malnutrition and it affects approximately 200 million children worldwide. Despite its importance and prevalence, it is not clear whether diminished growth affects brain development and neurocognitive outcomes. Further, diminished growth is more common in areas of extreme poverty, raising the possibility that it may serve as a mechanism for previously shown links between poverty and brain development. To address these questions, 79 children growing up in an extremely poor, urban area of Bangladesh underwent MRI at 6 years. Structural brain images were submitted to Mindboggle software, a Docker-compliant and highly reproducible tool for tissue segmentation and regional estimations of volume, surface area, cortical thickness, travel depth, and mean curvature. Diminished growth predicted brain morphometry and mediated the link between poverty and brain morphometry most consistently for white matter and subcortical volumes. Meanwhile, brain volume in left pallidum and right ventral diencephalon mediated the relationship between diminished growth and full-scale IQ. These findings offer malnutrition as one possible mechanism by which poverty affects brain development and neurocognitive outcomes in areas of extreme poverty.

## 1 Introduction

Early adverse experiences can substantially derail typical child development (1). This derailment is especially pronounced in communities of extreme poverty, where biological hazards such as malnutrition prevent children from reaching their full growth potential (2, 3), instead causing diminished growth (4–8), poor neurocognitive outcomes (5, 9–11), and premature death (4, 5, 12).

Globally, over 300 million children grow up in extreme poverty (UNICEF; https://www.unicef.org/social-policy/child-poverty). To monitor the impact of biological hazards on these children, many low-resource countries use the World Health Organization (WHO) anthropometric indicators: stunting, measured with height-for-age (HAZ); underweight, measured with weight-for-age (WAZ); and wasting, measured with weight-for-height (WHZ) (13). These are well-established proxies for malnutrition (4–8) that exhibit the greatest prevalence in areas with the highest rates of childhood poverty (2). Stunting, is the most common anthropometric indicator, occurring worldwide in over 150 million children under five (12), over 60 million of whom are from South Asia (2).

Diminished growth is also associated with reduced cognitive outcomes (5, 9–11, 14), leading to the hypothesis that it would also be associated with altered brain development (15). Indeed, greater stunting has been associated with less total white matter volume in infancy (16) and greater functional connectivity as measured with electroencephalography (EEG) in toddlerhood, which has been attributed to reduced pruning (11). These findings bolster earlier work describing cerebral atrophy in children with protein energy malnourishment (17–19), decreased intracranial volume in adults exposed to malnutrition prenatally (20), and white matter alterations in both (18, 20).

Importantly, malnutrition has a reciprocal relationship with inflammation— e.g., lack of key nutrients can engender immune dysfunction, infections, and inflammatory responses and sick children take in fewer nutrients due to lack of appetite (21). Consequently, when considering links between diminished growth and brain development it is also important to consider inflammation. Heightened inflammation, measured in blood or maternal amniotic fluid and in most cases attributed to infection, has been associated with reduced total intracranial volume (22) and white matter injury (23–25). Overall, these studies strongly point to susceptibility of brain structure, particularly white matter, to malnutrition. However, these findings were limited to presence/absence of brain injury, total tissue volumes, and/or relatively small sample sizes. To our knowledge, to date, no comprehensive examination of brain morphometry (e.g., with volumetric and surface-based measures of specific brain areas) in the context of malnutrition has been conducted in children.

In addition to malnutrition, a further consideration is the role of poverty in shaping brain development. In high-resource settings, socioeconomic status (SES) has been linked with total brain volume, total gray and white matter volumes, total surface area, and average cortical thickness (26). Regional hippocampus (26–30), amygdala (26, 27, 29), thalamus (26), and striatum (26) volumes were also associated with SES. In our previous work, we did not identify relations between measures of brain volume and measures of SES in settings of extreme poverty, defined by the World Bank as $1.90 per person per day. However, this work was conducted in infants without prolonged exposure to the multidimensional impacts of poverty, and it is unclear whether and, if so, how brain volume would relate to SES in children with prolonged exposure to these impacts.

Further, the mechanisms by which poverty alters brain development remain unclear. Poverty is thought to act indirectly through myriad risk factors, such as malnutrition and stress (15, 31). However, testing these mechanisms in formal mediation models is needed to confirm this hypothesis. Most studies examining these indirect effects have done so in the context of stress, finding that it mediates SES and hippocampal volume (29) and frontal cortex activation (32, 33). Family caregiving has also been shown to mediate SES and hippocampal volume (29). To our knowledge, no study has examined malnutrition as a potential mediator in the link between poverty and brain development, which represents a critical gap in the literature examining the effect of SES on child’s outcomes. In addition, as with direct associations with poverty, it is unclear whether these indirect effects would be the same or different in children growing up in extreme poverty.

The present study addresses three main lines of inquiry. The first line is whether and how early diminished growth predicts subsequent brain development, specifically morphometry. To examine this, we collected anthropometry at age 2 years and structural MRI at 6 years in 79 children growing up in an extremely poor, urban area of Bangladesh, a vastly underrepresented population in neuroscientific research. To comprehensively examine brain morphometry, MRI data were submitted to Mindboggle (34), a software that estimates volume, surface area, cortical thickness, travel depth, and mean curvature in a Docker container for high reproducibility (35). We hypothesized associations between diminished growth and brain volume, surface area, and cortical thickness based on common links between these measures and SES (2, 26). We did not expect associations between diminished growth and travel depth or mean curvature because the latter two are thought to be primarily under genetic control (36), suggesting less susceptibility to environmental factors such as malnutrition.

The second line of inquiry concerns the broader context of poverty in altering brain development. Particularly, we examined whether poverty is associated with brain morphometry as was reported for high-resource settings (26–28) and whether malnutrition mediates an association between poverty and brain morphometry. In our previous work, poverty was not linked with diminished growth or brain volume in infancy (16), but in our previous report these two measures were acquired in infancy, before children had prolonged exposure to poverty. That the relationship between diminished growth and poverty is age-dependent (37), with links at 3 years but not in infancy (38), suggests that poverty would now relate to diminished growth and brain morphometry in our cohort of 6 year-olds. The third line of inquiry addresses whether brain morphometry mediates associations between diminished growth and neurocognitive outcomes.

## 2 Methods

### 2.1 Participants

The present study is part of the Bangladesh Early Adversity Neuroimaging (BEAN) study (10, 11, 16, 38–41), investigating effects of extreme poverty on early brain development in children in Dhaka, Bangladesh. The overall study followed 260 children beginning in infancy, from whom socioeconomic, anthropometric, behavioral, and biological measures were collected. The current study focused on the 81 children who underwent structural MRI between ages six and eight. After excluding children with severe motion artifacts, the final sample comprised 79 structural MRI datasets (6.68 ± 0.40 years; F/M = 36/43; please see Table 1 for details). No child had been diagnosed with a neurological disease or disability and all scans were reviewed for malignant brain features by a clinical radiologist in Bangladesh and a pediatric neuroradiologist at Boston Children’s Hospital. The study was approved by research and ethics review committees at BCH and The International Centre for Diarrhoeal Disease Research, Banglades

**Table 1.** Subject Demographics

| Table 1. Subject Demographics |  |
| --- | --- |
| N | 79 |
| Sex (F/M) | 36/43 |
| Age at MRI (Years) | $6.7 \pm 0.40$ |
| Age Range at MRI (Years) | 5.5 - 7.0 |
| HAZ<sup>a</sup> | $-1.6 \pm 0.89$ |
| WAZ<sup>a</sup> | $-1.3 \pm 1.1$ |
| WHZ<sup>a</sup> | $-0.66 \pm 1.1$ |
| Maternal Education (Years)<sup>b</sup> | $4.4 \pm 3.9$ |
| Income-to-Needs<sup>c</sup> | $3100 \pm 1700$ |
| Full-Scale IQ | $86 \pm 9.1$ |
| <sup>a</sup>Scores less than -2 indicate stunting, underweight, or wasting (2) |  |
| <sup>b</sup>Rescaled (please see Methods) |  |
| <sup>c</sup>Monthly family income in Bangladeshi Taka/number of household members |  |

### 2.2 Anthropometric Measures

Height-for-age (HAZ), weight-for-age (WAZ), and weight-for-height (WHZ) scores were used to estimate stunting, underweight, and wasting, respectively. Trained, local staff measured height (in centimeters) and weight (in kilograms). These measures were then submitted to the Anthro Plus software (https://www.who.int/tools/child-growth-standards/software; Multicenter Growth Reference Study (42)) and compared with growth curve data from 8440 infants (0–24 months) and children (18–71 months) from Brazil, Ghana, India, Norway, Oman and the U.S. Resulting z-scores (i.e., HAZ, WAZ, and WHZ), which were age- and sex-referenced and standardized, reflected deviations from typical growth trajectories. Critically, the standard growth curves comprised infants and children who grew up in healthy environments (including with breastfeeding and absence of smoking) that are “likely to favour the achievement of their full genetic growth potential” (43). As such, consistent deviations from these standard growth curves can be inferred to reflect environmental hazards during upbringing. Lastly, anthropometry was assessed at 21, 30, and 36 months and these values were averaged to ensure stability of these measures. Stunting, under-weight, and wasting were respectively defined as HAZ, WAZ, and WHZ less than -2 (2). According to this definition, 24 children were stunted, 18 children were underweight, and 8 children were wasted. Six children were stunted, underweight, and wasted. Table 1 summarizes these measures in the final cohort.

### 2.3 Measures of Poverty

Years of maternal education, monthly family income, and number of household members were used to compute two measures of poverty: maternal education and income-to-needs ratio. Maternal education was measured as years of formal education, ranging from 0 to 10, with 0 indicating no formal education, 1–9 indicating number of grades passed, and 10 indicating education beyond the 9th grade passed, in which degrees may be conferred. Income-to-needs was computed as the monthly family income divided by the number of household members. As the exchange rate for USD to Bangladeshi taka is USD$1:Tk85, the family in the cohort with the lowest monthly income-to-needs earned Tk890 or USD$10 per household member per month, while the family in the cohort with the highest monthly income-to-needs earned Tk10,000 or USD$120 per household member per month. When considering that the World Bank (https://data.worldbank.org/) international standard for extreme poverty is roughly Tk4800 or USD$57 per person per month (calculated from USD$1.90/day and assuming 30 days/month), the final cohort constituted 71 out of 79 children living in extreme poverty. Maternal education and income-to-needs were assessed twice—once at age 6 months and a second time at age 3 years—through oral interviews with the children’s parents, and then averaged, to better capture overall measures of poverty that reflected the entire childhood. Due to a positive skew in the income-to-needs variable, these data were log (base 10) transformed.

### 2.4 Neurocognitive Assessment

Children underwent neurocognitive testing using the Wechsler Preschool and Primary Scale of Intelligence (WPPSI) administered by trained, local psychologists and staff. Although the WPPSI has been standardized using U.S. children, steps were taken to ensure translatability in this Bangladeshi cohort. For instance, items in the assessment were translated and culturally adapted (38) and the Bangladeshi version of the assessment exhibited test-retest reliability (44, 45). The full-scale intelligence quotient (FIQ) score, which reflects children’s general cognitive abilities, was tested for associations with anthropometric and brain morphometric measures. While WPPSI was assessed at age five years, prior to MRI scanning (conducted between ages 5.5 and 7.0 years), these tests are considered relatively stable after four years of age (46).

### 2.5 MRI data acquisition

Neuroimaging data were acquired on a 3T Siemens MAGNETOM Verio scanner at the National Institute for Neuroscience and Hospital (NINSH) in Dhaka, Bangladesh. Consenting was done at NINSH on the day prior to scanning. For consenting and scanning, children and their mothers were brought to and from NINSH by study staff, usually via rickshaw. The cost of transport was paid for by the study and children and mothers were provided meals on the day of the scan.

NINSH previously scanned pediatric patients for clinical examinations and for pilot studies in infants (16, 41), but this study marks the first time this facility had collected MRI data for a large-scale pediatric neuroimaging study. As such, local staff visited Boston Children’s Hospital to receive training on conducting pediatric MRI studies (47). Subsequently, a protocol was designed that incorporated this training and the limitations of a low-resource setting (for a general description of challenges of conducting MRI in a low resource setting, please see (41)). specifically, children were brought to NINSH by trained, local staff. Prior to scanning, children were shown sample brain structural images to teach them about the purpose of their visit and to explain that the machine they would enter would take pictures of their brain. They then went through several steps to practice remaining motionless. First, they were shown MRI images from a child remaining motionless during scanning (i.e., a clear image) and a child moving during scanning (i.e., a blurry image) to understand the effects of motion. Second, staff played a ”freeze tag” game with the them, such that children practiced becoming still when staff instructed them to ”freeze.” Third, they were asked to lie motionless for 1 minute and offered encouragement for doing so. If they moved while doing this, they were told that any tapping they felt on their feet meant that they need to remain still.

Following this training, children saw and entered a cardboard mock scanner to familiarize them with the scanning environment (Fig. 1). Staff narrated throughout to ensure children would not be scared. Children again received a tapping on their feet to practice lying motionless. To ensure children would become familiar with scanner noise, staff tapped on the outside of the cardboard box and played MRI sounds previously recorded on a CD. Children were then shown pictures of the real scanner so that they could connect its makeup to that of the mock scanner.

**Figure 1.**
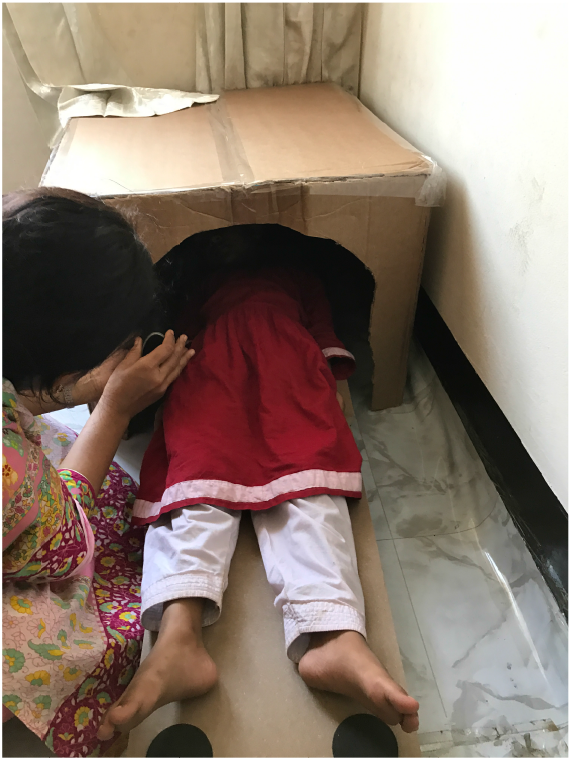
Cardboard mock scanner at MRI facility.

Structural T1-weighted magnetization-prepared rapid gradient-echo (MPRAGE) scans were acquired with the following parameters: TR = 2500 ms, TE = 3.47 ms, 176 sagittal slices, 1 mm3 voxels, FOV = 256 mm. Functional and diffusion scans were also acquired but will be described in future reports.

### 2.6 MRI data processing

Images were visually inspected for artifacts. Following the removal of two artifactual datasets, the remaining raw MPRAGE images were processed using Mindboggle 1.3.8 and run in a Docker container (https://Mindboggle.readthedocs.io/en/latest/ (34)). This pipeline implements Advanced Normalization Tools (ANTs) and FreeSurfer (v6.0.0). First, Mindboggle calls antsCorticalThickness, which includes brain extraction, N4 bias correction, tissue segmentation, and cortical thickness estimation. Subsequently, Mindboggle submits raw images to recon-all, which segments the brain into different tissue classes, approximates pial surfaces, and labels volumes and surfaces by brain region.

The final series of steps belong to Mindboggle proper. First, FreeSurfer output is converted to nifti and vtk filetypes for combining with ANTs. Second, Mindboggle runs a hybrid segmentation algorithm to reconcile differences between ANTs and FreeSurfer segmentations. These toolkits separately mislabel tissue classes in different ways, with ANTs underestimating white matter and including non-brain tissue, while FreeSurfer omits gray matter voxels by over-cropping. Third, volumetric measures are computed for cortical and subcortical brain regions (including white matter), while surface area, cortical thickness (from FreeSurfer), travel depth, and mean curvature measures are computed for each cortical brain region. Two sets of measures were provided for brain volume of various structures—one using ANTs labels and another using FreeSurfer labels. We opted to use the latter for two reasons. First, cortical thickness measures were reported as FreeSurfer-derived. Second, FreeSurfer’s labels were more comprehensive for white matter by comparison with ANTs and white matter has been particularly relevant in the context of anthropometry (16). Mindboggle also offered geodesic depth; however, we limited our analyses to travel depth as these two measures are highly similar across the brain except in insular regions (34)), where we did not have specific hypotheses. Laplace-Beltrami spectrum and Zernicke moments output by Mindboggle were also excluded from analyses. As with other neuroimaging tools that run via the Docker (e.g., fMRIPrep (48), this pipeline is highly reproducible.

In total, we examined global measures of brain volume (total brain volume, total gray matter volume, total white matter volume) and global surface-based measures (total surface area, average cortical thickness, average travel depth, and average mean curvature). Mindboggle outputs these brain measures by region. This made computations of global estimates for volume and surface area (for which summing would be employed) straightforward, but hampered global estimations for cortical thickness, travel depth, and mean curvature (for which summing would have been difficult to interpret), because these regions were of varying sizes. To circumnavigate this challenge, we computed average travel depth, average curvature, and average cortical thickness weighted by surface area in each region. We then examined regional measures of brain volume from cortical, subcortical, and white matter areas, as well as regional surface measures (surface area, cortical thickness, travel depth, and curvature). Lastly, one volumetric estimate was by default labelled as ‘unsegmented white matter.’ Upon visual inspection of this region, the vast majority of voxels iwere in corona radiata and internal capsule. Given this, in our reporting, we labelled ‘unsegmented white matter’ as ‘corona radiata and internal capsule.

### 2.7 Statistical Analyses

Previous studies by our group have reported on associations between measures of poverty and anthropometry (38). We also examined this relationship in this cohort of 6 year-olds by testing for correlations between anthropometric measures and maternal education and (log of) income-to-needs.

To address our first line of inquiry, that diminished growth predicts brain morphometry, we submitted total and regional volumetric and surface-based measures to semipartial correlation analyses controlling for age at time of scan and sex. False discovery rate (FDR) corrections for multiple comparisons were applied separately for volumetric, surface area, depth, curvature, and thickness measures and for total and regional analyses (49). However, tests with HAZ, WAZ, and WHZ were corrected for multiple comparisons altogether. To address whether poverty predicts brain morphometry, as hypothesized, this procedure was repeated replacing anthropometric measures with measures of maternal education and income-to-needs.

We also hypothesized that diminished growth mediates relationships between measures of poverty and brain morphometry. To test this, indirect effects were examined whenever an anthropometric measure exhibited a significant (after FDR correction for multiple comparisons) association with measures of poverty and brain morphometry. Indirect effects were reported as significant when 95% confidence intervals (based on 10,000 bootstrapped samples) did not include 0 and as proportion of the total effect (indirect + direct effect = total effect). This process was repeated for our third line of inquiry: whether measures of brain morphometry mediated associations between anthropometric measures and neurocognitive outcomes (i.e., FIQ).

Correlational tests were conducted in Matlab and indirect effects were examined using the Mediation package in R. All brain maps were generated using the ggseg() function in R.

## 3 Results

### 3.1 Poverty is associated with diminished growth

We first examined the association between anthropometric indicators of stunting (i.e., HAZ), underweight (i.e., WAZ), and wasting (i.e., WHZ) and measures associated with poverty. Maternal education was positively related to HAZ (r = 0.29, p = 0.0088), WAZ (r = 0.27, p = 0.017), and WHZ (r = 0.20, p = 0.083). Income-to-needs, which was log-transformed due to its skew (28), was associated with HAZ (r = 0.27, p = 0.017), WAZ (r = 0.30, p = 0.0075), and WHZ (r = 0.26, p = 0.021). All were significant at p_FDR_ *<*0.05 except for between maternal education and WHZ. In summary, we observed that measures of poverty were associated with diminished growth.

### 3.2 Diminished growth predicts brain morphometry

We next examined the association between anthropometric measures for diminished growth and measures of brain structure, including global and regional volumetric and surface-based measures. Global measures of brain morphometry are summarized in Table 2. In terms of volumetric measures, total brain volume (TBV), total gray matter volume (GMV), and total white matter volume (WMV) were positively associated with HAZ, WAZ, and WHZ (p_FDR_ *<*0.05; Fig. 2). Total surface area (SA) across all cortical regions was associated with HAZ and WAZ (p_FDR_ *<*0.05), but not WHZ (p_unc_ *>*0.05). Average travel depth was also positively associated with HAZ (pFDR ¡ 0.05), but not WAZ or WHZ (p_unc_ *>*0.05). Neither average cortical thickness nor average mean curvature was associated with any anthropometric measures (p_unc_ *>*0.05).

**Table 2.** Global Estimates of Brain Morphometry

| Table 2. Global Estimates of Brain Morphometry |  |
| --- | --- |
| Total brain volume (mm<sup>3</sup>) | 1087300 (82428) |
| Total gray matter volume (mm<sup>3</sup>) | 685850 (55271) |
| Total white matter volume (mm<sup>3</sup>) | 374940 (32442) |
| Total surface area (mm<sup>2</sup>) | 199000 (17371) |
| Average Cortical Thickness (mm) | 2.9092 (0.0995) |
| Average Travel Depth (mm) | 5.9399 (0.3024) |
| Average Curvature (mm) | -3.6173 (0.4230) |

**Figure 2.**
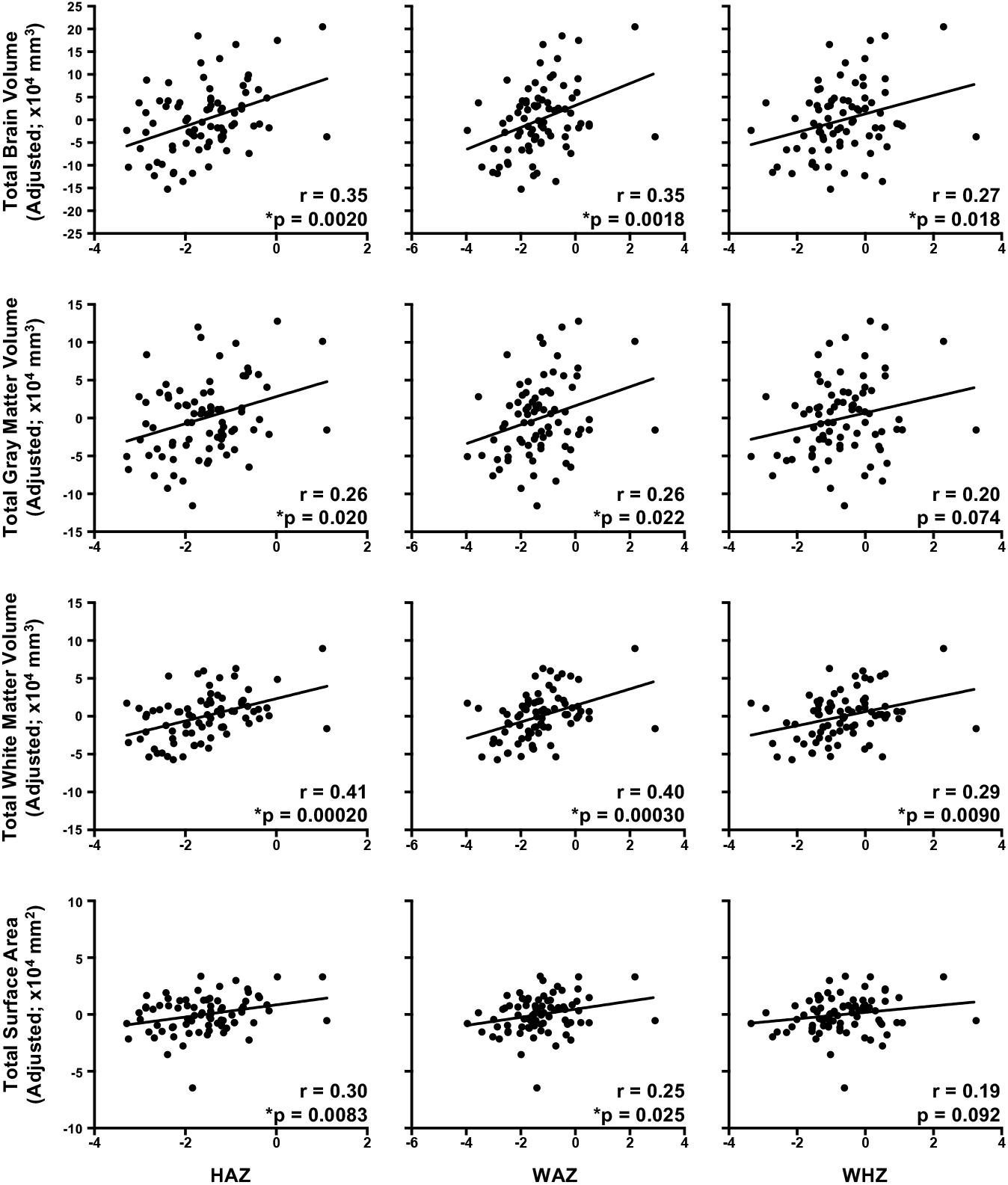
Diminished growth measured at 2 years predicts global measures of brain morphometry. Relationships were computed using semipartial correlations with brain measures adjusted for age and sex. Strongest effects were observed for total white matter volume and for HAZ. Please note, FDR correction for multiple comparisons were performed separately for volumetric and surface-based measures. N = 79. *p_FDR_ *<*0.05.

To investigate whether global measures of volume, surface area, and travel depth were driven by specific brain regions, we also examined brain-anthropometric associations in all brain regions delineated by Mindboggle. Volumetric measures were positively associated with HAZ and WAZ mainly in subcortical gray matter and white matter regions, and with WHZ only in white matter regions (Fig. 3; Supplementary Table 3). Surface area in right caudal anterior cingulate was associated with HAZ (r = 0.39; p = 0.00044). However, no other surface-based measures were associated with anthropometry. Overall, diminished growth predicted brain morphometry, and these associations were most robust between HAZ and WAZ and brain volume in subcortical and white matter regions.

**Figure 3.**
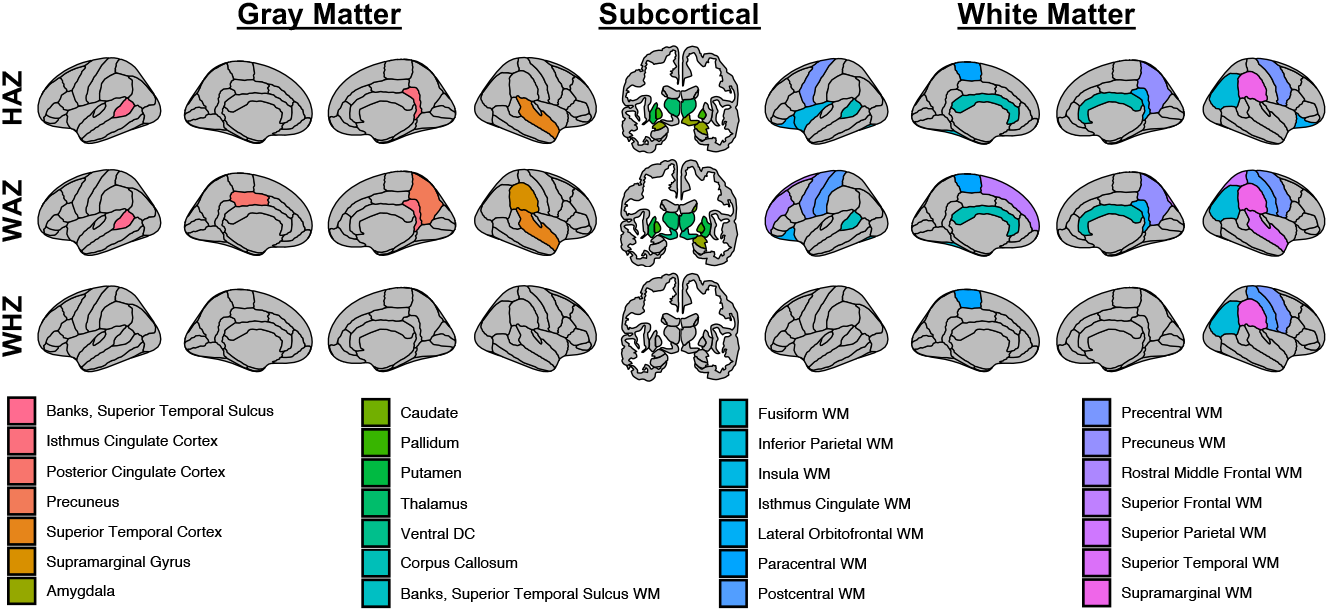
Diminished growth predicts regional brain volume. Associations occur mostly in white matter and subcortical regions and for HAZ and WAZ. All brain maps p_FDR_ *<*0.05 FDR corrected for multiple comparisons for each anthropometric indicator. Please note that corona radiata and internal capsule are not depicted.

### 3.3 Poverty is associated with diminished growth

We next examined whether global and regional volumetric and surface-based measures were associated with measures of poverty. In terms of global measures, TBV, GMV, WMV, and SA were positively associated with maternal education and income-to-needs (p_FDR_ *<*0.05; Fig. 4), which was consistent with brain-anthropometry findings. No other surface-based measures were associated with either maternal education nor income-to-needs.

**Figure 4.**
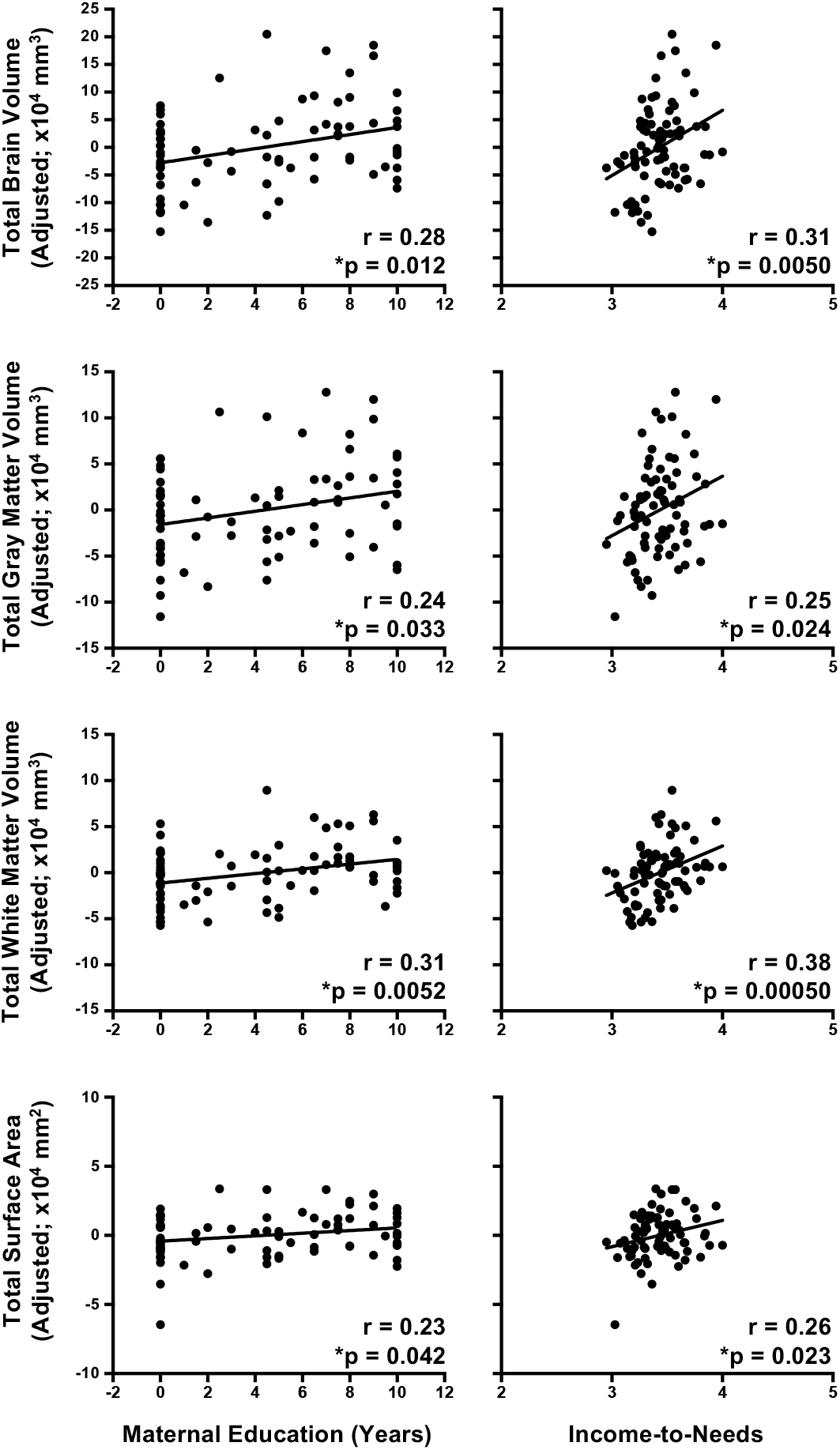
Childhood poverty as measured with maternal education and income-to-needs predicts global measures of brain morphometry. Relationships were computed using semipartial correlations with brain measures adjusted for age and sex. Income-to-Needs is computed as the log of household monthly income in Bangladeshi Taka divided by number of household members. Strongest effects were observed for total white matter volume. Please note, FDR correction for multiple comparisons were performed separately for volumetric and surface-based measures. N = 79. *p_FDR_ *<*0.05.

Subsequently, regional associations between measures of poverty and brain morphometry were examined. As with anthropometric measures, volumetric measures were associated with maternal education and, to a lesser extent, income-to-needs mainly in subcortical gray matter and white matter regions (Supplementary Table 2). Nevertheless, the only direct overlap of brain regions exhibiting brain-poverty and brain-anthropometry associations were in bilateral pallidum, putamen, ventral diencephalon, and corona radiata and internal capsule; left thalamus, fusiform white matter, and lateral orbitofrontal white matter; and right isthmus cingulate cortex. The only surface-based measure associated with measures of poverty was with surface area, namely, between maternal education and right pars triangularis (r = 0.35; p = 0.0016; p_FDR_ *<*0.05) and between maternal education and right rostral middle frontal cortex (r = 0.36; p = 0.00097; p_FDR_ *<*0.05).

### 3.4 Diminished growth mediates the relationship between poverty and brain morphometry

We next examined indirect pathways between poverty and brain morphometry via diminished growth. As a precondition for mediation, the mediator (i.e., diminished growth) must be associated with both the predictor (i.e., measure of poverty) and outcome (i.e., brain morphometry). Thus, we examined indirect effects only where HAZ, WAZ, or WHZ were related to maternal education or income-to-needs and volumetric or surface-based measures. Further, indirect effects are only reported where 95% confidence intervals (CI), based on 10,000 bootstrapped samples, do not include 0.

For global volumetric measures, HAZ mediated the relationship between maternal education and TBV (proportion mediated = 0.32, CI [0.044 0.89], p = 0.018) and TWM (proportion mediated = 0.36, CI [0.074 1.04], p = 0.0094) and between income-to-needs and TBV (proportion mediated = 0.29, CI [0.046 0.81], p = 0.015) and TWM (proportion mediated = 0.29, CI [0.067 0.68], p = 0.0064). WAZ also partially mediated the association between maternal education and TBV (proportion mediated = 0.27, CI [0.015 0.82], p = 0.034) and TWM (proportion mediated = 0.31, CI [0.034 0.94], p = 0.021) and between income-to-needs and TBV (proportion mediated = 0.28, CI [0.012 0.78], p = 0.040) and TWM (proportion mediated = 0.30, CI [0.048 0.70], p = 0.015). No indirect effects were observed for WHZ. Additionally, associations between measures of poverty and surface-based brain measures were not mediated by diminished growth.

We next examined indirect effects with regional brain measures. Unlike with global brain measures, which were correlated with both predictor and mediator variables, very few regional measures were correlated with both predictors and mediators (bilateral pallidum, putamen, ventral diencephalon, and corona radiata and internal capsule; left thalamus, fusiform white matter, and lateral orbitofrontal white matter; and right isthmus cingulate cortex). However, modern models of mediation do not stipulate an association between predictor and outcome (50). Therefore, we examined indirect pathways for any brain region associated with an anthropometric measure. As with associations between diminished growth and brain volume, indirect effects were observed mainly for subcortical and white matter regions (Fig. 5). Notably, relatively consistent mediations (across several anthropometric and poverty measure pathways; please see Supplementary Table 3) were observed for right amygdala, corpus callosum, and bilateral corona radiata and internal capsule. In summary, diminished growth mediated the relation between maternal education and income-to-needs and global and regional (mainly subcortical and white matter) volumes.

**Figure 5.**
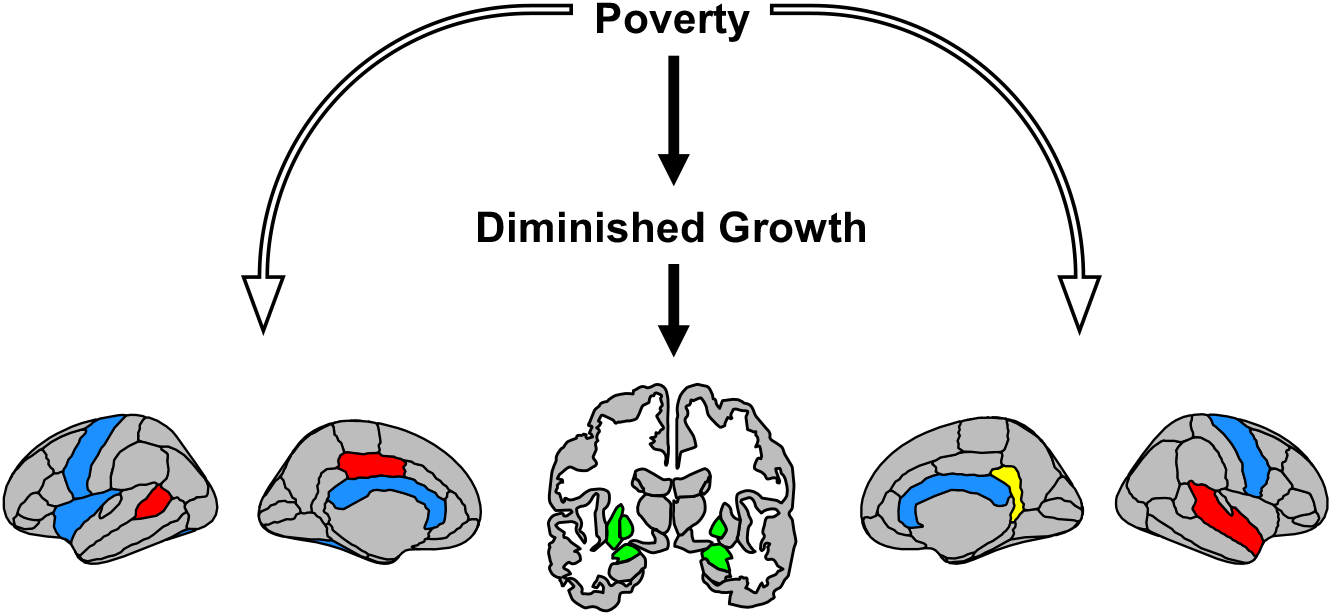
Diminished growth mediates the relationship between poverty and regional brain volume. Poverty measures include maternal education and income-to-needs. Diminished growth includes HAZ, WAZ, and WHZ. Brain maps show all brain regions whose association with measures of poverty are mediated by diminished growth. Indirect effects (filled arrows) were in mainly in white matter (blue) and subcortical gray matter (green), with some exceptions in gray matter (red) and right isthmus cingulate (yellow), for which white and gray matter exhibited indirect effects. Direct pathways (unfilled arrows) are also shown to reflect that diminished growth mediates a proportion, but not all, of the association between poverty and brain volume. Please note that corona radiata and internal capsule are not depicted.

### 3.5 Brain morphometry mediates the association between diminished growth and IQ

We next examined relationships between anthropometric measures and full-scale IQ (FIQ) scores. Full-scale IQ was predicted by HAZ (r = 0.35, p = 0.0016, p_FDR_ *<*0.05), WAZ (r = 0.37, p = 0.00080, p_FDR_ *<*0.05), and WHZ (r = 0.33, p = 0.0034, p_FDR_ *<*0.05). Global measures of brain morphometry were also related to FIQ, namely TBV (r = 0.28, p = 0.012), GMV (r = 0.27, p = 0.017, p_FDR_ *<*0.05), average cortical thickness (r = 0.26, p = 0.022, p_FDR_ *<*0.05), and average travel depth (r = 0.25, p = 0.0257), as were regional volumes in left pallidum (r = 0.45, p = 0.000027, p_FDR_ *<*0.05), and right ventral diencephalon (r = 0.42, p = 0.00010, p_FDR_ *<*0.05).

Finally, we tested indirect effects between diminished growth and FIQ via global and regional measures of brain morphometry for the ten brain measures correlated with both anthropometry and FIQ. Of these, mediation effects were observed for the pathways between HAZ and FIQ via left pallidum (proportion mediated = 0.39, CI [0.11 0.99], p = 0.0034) and right ventral diencephalon (proportion mediated = 0.30, CI [0.050 0.87], p = 0.013) and between WAZ and FIQ via left pallidum (proportion mediated = 0.35, CI [0.10 0.86], p = 0.0032) and right ventral diencephalon (proportion mediated = 0.29, CI [0.046 0.79], p = 0.015; Fig. 6). In summary, we observed links between diminished growth and FIQ and these relations were partially mediated by left pallidum and right ventral diencephalon volumes.

**Figure 6.**
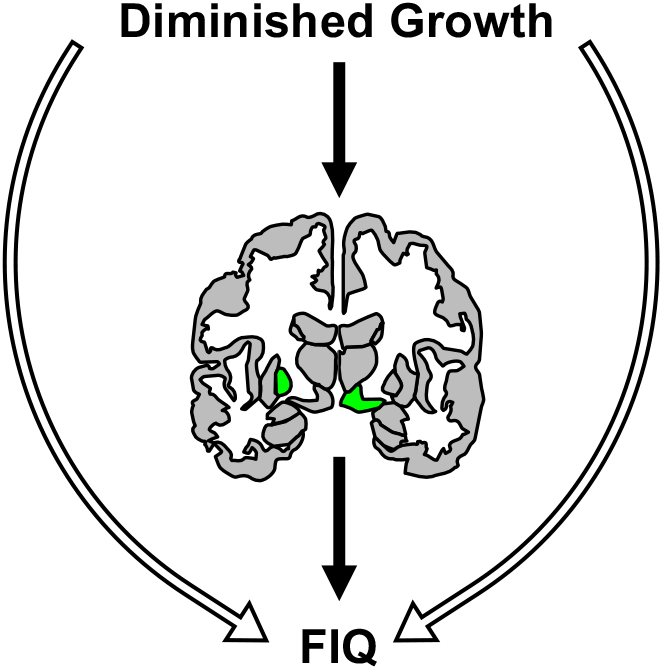
Brain volumes in left pallidum and right ventral diencephalon partially mediate the association between diminished growth and FIQ. Diminished growth includes HAZ and WAZ. Indirect pathways (filled arrows) show diminished growth to FIQ via brain volume and direct pathways (unfilled arrows) clarify that left pallidum and right ventral diencephalon volumes mediate a proportion, but not the entirety, of the association between diminished growth and IQ.

## 4 Discussion

Malnutrition occurs frequently in children growing up in extreme poverty and is thought to substantially derail typical cognitive development (15). In this study, we observed that diminished growth—i.e., stunting, underweight, and wasting—which are proxies for malnutrition, strongly predicted brain morphometry and these effects were strongest in subcortical and white matter regions. Further, diminished growth mediated links between measures of childhood poverty and brain volume. Finally, left pallidum and right ventral diencephalon volumes partially mediated the associations between diminished growth and full-scale IQ (FIQ).

Diminished growth at roughly 2 years predicted global and regional morphometric measures at 6 years, including total brain volume, total gray matter volume, total white matter volume, and total surface area. As predicted, findings overall do not support robust relationships between diminished growth and travel depth or mean curvature. The particularly strong predictions of global white matter volume from diminished growth supports earlier work by our group showing that stunting and underweight inversely relate to total white, but not total gray, matter volume in infancy (16). Results are also consistent with clinical findings of gross white matter atypicalities in children suffering from protein energy malnutrition (18, 20) and inflammation (23–25), which is strongly linked with malnutrition (21). The relationship between inflammation and white matter has a basis in animal work, with inflammation disrupting oligodendrocyte maturation and consequently reducing myelination (51). We add to this literature with quantitative estimates of white matter volume and with topographical maps that depict where diminished growth predicts white matter volume.

With regard to subcortical structures, findings in the ventral diencephalon, which includes the hypothalamus (52), were also compelling because this area is thought to be a nexus for complex brain-gut-inflammation interactions (53). specifically, we identified an association between brain volume and a measure that is thought to reflect malnutrition (to which inflammation is linked) in an area with bidirectional connections with the gut (54) and with a role in inflammatory regulation via the hypothalamic-pituitary-adrenal (HPA) axis (55). Moreover, the gut microbiome is thought to play a vital role in the development of the HPA axis (56). That ventral diencephalon was shown to mediate the relationship between diminished growth and FIQ further supports this, as well as the hypothesis that this area affects cognitive function (57). Left pallidum was also shown to mediate the relationship between diminished growth and IQ, and this area may affect cognition through known anatomical projections to prefrontal cortex, as shown in non-human primates (58).

Our work also addresses a persistent gap in neuroscientific research on poverty, namely that the overwhelming majority of such research has been conducted in high-resource settings. Our findings in children growing up in a low-resource setting support studies in high-resource countries reporting associations between socioeconomic status (SES) and total brain volume (26), gray matter volume (26, 29), white matter volume (26, 29), and surface area (26, 28). Regional effects were observed in the striatum, which was also found previously (26). The most notable departures from studies in high-resource settings were the absences of effects for cortical thickness (26, 28) and for the hippocampus (26–28, 59), suggesting that the brain correlates of extreme poverty may differ from the correlates of SES in high-resource countries. Additionally, by comparison with reports of age-comparable, typically developing children growing up in the U.S., the cohort in Bangladesh exhibited lower total brain volume (60) and total gray and white matter volume (61, 62). Future cross-cultural studies will need to directly compare brain volumes processed using identical methods to confirm this.

Our findings also speak to another consistent gap in the SES literature, which relates SES to brain structure and function, but so far has not identified many intermediary hazards that poverty might work through to effect brain alterations (for a discussion, please see (31)). As hypothesized, we found that diminished growth mediated the relationship between poverty and global and regional brain volume. It is noteworthy that links between both measures of poverty—maternal education and income-to-needs—and right amygdala were mediated by two of the three measures of diminished growth—HAZ and WAZ. This region’s volume exhibited associations with SES in some (26, 29, 41), but not other SES studies (28, 30, 63), and it was proposed that the presence of an SES-amygdala association may depend on the degree of poverty experienced by the study cohort (64), which is consistent with our findings in children growing up in extreme poverty.

The links of poverty to diminished growth and to brain morphometry via indirect and direct pathways merits discussion in the context of our earlier work, in which we did not observe these links. Unlike the present study, the prior work was conducted in children at infancy, before long-term exposure to poverty (16), suggesting that the effects of poverty take time to accumulate (65). This explanation also makes sense when considering that the association between diminished growth and poverty is age-dependent (37), with poverty linked with anthropometry at 3 years, but not in infancy (38). Overall, measures of poverty have widespread and substantial associations with subcortical and white matter volumes and these associations are partially mediated by diminished growth (as a proxy of malnutrition).

The present work had three notable limitations. The first is that diminished growth is a proxy for malnutrition (4–8), rather than a direct measure. However, malnutrition, for instance, is a complex factor, and using specific measures of nutrient deficiencies (e.g., zinc or iron) may not have adequately captured the full impact of myriad deficiencies on human development. A second limitation is that HAZ, WAZ, and WHZ were all examined as potential mediators. While each were involved in significant indirect pathways separately (after FDR correction), their independent contributions as mediators is complicated by collinearity between them (e.g., HAZ and WAZ: r = 0.87; p = 3.4 x 10-25). For this reason, indirect pathways linking poverty to brain morphometry are characterized as mediated by diminished growth in general. A final limitation is that the measures for diminished growth and poverty were not contemporaneous with MRI acquisition. However, while it is possible that these measures would change by the time the child reaches six years, at least across measurements, these estimates were relatively constant (e.g., HAZ at 21 and 36 months were correlated at r = 0.91; p = 6.6 x 10-50 in a larger sample of children).

Finally, the vast majority of databases comprise data from white U.S. and European individuals (66). In contrast, the children in this study grew up in an impoverished area of Bangladesh, making them an extremely underrepresented population in neuroscientific research, both racially and socioeconomically. As such, involving them in MRI studies such as the present one is critical to begin to address inequity in developmental cognitive neuroscience research. We encourage others to chart similar paths in their own lines of inquiry.

## Conclusion

This study comprehensively examines links among brain morphometry, diminished growth, poverty, and neurocognitive outcomes in children growing up in a low-resource setting in Bangladesh. Our findings show that diminished growth, a proxy for malnutrition, predicts brain morphometry and mediates associations between poverty and brain morphometry. In doing so, this study implicates malnutrition as one pathway by which poverty may alter brain development. Although future longitudinal studies with nutritional interventions will be needed to test the causality of this pathway, this study has important implications for the role of nutrition in children growing up in low-resource settings.

## 5 Acknowledgements

This study was funded by research grants from the Bill Melinda Gates Foundation to CA Nelson [OPP1111625] and WA Petri. [OPP1017093], and research grants to WA Petri from the Henske Foundation and the NIAID (R01 AI043596-17). We are grateful to the families who participated in the study, the staff at The International Centre for Diarrhoeal Disease Research who undertook and completed data collection, and Uma Nayak for organizing the database of non-MRI measures. Finally, we thank the Harvard Catalyst Biostatistical Consulting program for guidance on reporting mediations.

---

**Supplementary Table 1.** Brain-Anthropometry Relationships

| *HAZ* | **r** | **p (x 10<sup>-3</sup>)** |
| --- | --- | --- |
| L. Banks, Superior Temporal Sulcus | 0.31 | 4.9 |
| R. Isthmus Cingulate Cortex | 0.33 | 3.0 |
| R. Superior Temporal Cortex | 0.32 | 3.7 |
| L. Amygdala | 0.31 | 5.9 |
| L. Cerebellum White Matter | 0.38 | 0.55 |
| L. Pallidum | 0.43 | 0.063 |
| L. Putamen | 0.32 | 3.9 |
| L. Thalamus | 0.32 | 3.8 |
| R. Amygdala | 0.39 | 0.39 |
| R. Cerebellum White Matter | 0.36 | 1.1 |
| R. Pallidum | 0.39 | 0.40 |
| R. Thalamus | 0.30 | 6.8 |
| R. Ventral Diencephalon | 0.33 | 3.1 |
| Corpus Callosum | 0.33 | 2.9 |
| L. Banks, Superior Temporal Sulcus White Matter | 0.30 | 6.6 |
| L. Fusiform White Matter | 0.45 | 0.039 |
| L. Insula White Matter | 0.31 | 4.7 |
| L. Lateral Orbitofrontal White Matter | 0.34 | 2.0 |
| L. Paracentral White Matter | 0.31 | 5.0 |
| L. Precentral White Matter | 0.30 | 6.4 |
| L. Corona Radiata and Internal Capsule | 0.39 | 0.40 |
| R. Inferior Parietal White Matter | 0.44 | 0.061 |
| R. Isthmus Cingulate White Matter | 0.34 | 2.0 |
| R. Lateral Orbitofrontal White Matter | 0.32 | 4.3 |
| R. Precentral White Matter | 0.31 | 5.3 |
| R. Precuneus White Matter | 0.33 | 2.6 |
| R. Supramarginal White Matter | 0.32 | 4.6 |
| R. Corona Radiata and Internal Capsule | 0.44 | 0.045 |
| *WAZ* |  |  |
| L. Banks, Superior Temporal Sulcus | 0.30 | 8.1 |
| L. Posterior Cingulate Cortex | 0.30 | 7.0 |
| R. Isthmus Cingulate Cortex | 0.32 | 4.4 |
| R. Precuneus | 0.29 | 9.5 |
| R. Superior Temporal Cortex | 0.33 | 3.0 |
| R. Supramarginal Cortex | 0.31 | 4.9 |
| L. Cerebellum White Matter | 0.39 | 0.41 |
| L. Pallidum | 0.44 | 0.052 |
| L. Putamen | 0.29 | 10 |
| L. Thalamus | 0.35 | 1.4 |
| L. Ventral Diencephalon | 0.33 | 3.0 |
| R. Amygdala | 0.34 | 2.2 |
| R. Caudate | 0.30 | 7.4 |
| R. Cerebellum White Matter | 0.33 | 2.6 |
| R. Pallidum | 0.37 | 0.94 |
| R. Putamen | 0.31 | 5.8 |
| R. Thalamus | 0.35 | 1.5 |
| R. Ventral Diencephalon | 0.38 | 0.62 |
| Corpus Callosum | 0.36 | 1.3 |
| L. Banks, Superior Temporal Sulcus White Matter | 0.34 | 2.3 |
| L. Fusiform White Matter | 0.38 | 0.47 |
| L. Lateral Orbitofrontal White Matter | 0.34 | 2.5 |
| L. Paracentral White Matter | 0.40 | 0.24 |
| L. Postcentral White Matter | 0.33 | 2.8 |
| L. Precentral White Matter | 0.31 | 4.7 |
| L. Rostral Middle Frontal White Matter | 0.29 | 8.3 |
| L. Superior Frontal White Matter | 0.29 | 10 |
| L. Corona Radiata and Internal Capsule | 0.44 | 0.052 |
| R. Inferior Parietal White Matter | 0.44 | 0.044 |
| R. Isthmus Cingulate White Matter | 0.35 | 1.7 |
| R. Postcentral White Matter | 0.34 | 2.2 |
| R. Precentral White Matter | 0.40 | 0.28 |
| R. Precuneus White Matter | 0.34 | 2.2 |
| R. Superior Parietal White Matter | 0.29 | 11 |
| R. Superior Temporal White Matter | 0.35 | 1.8 |
| R. Supramarginal White Matter | 0.41 | 0.16 |
| R. Corona Radiata and Internal Capsule | 0.50 | 0.0025 |
| *WHZ* |  |  |
| L. Paracentral White Matter | 0.36 | 1.1 |
| L. Corona Radiata and Internal Capsule | 0.38 | 0.54 |
| R. Inferior Parietal White Matter | 0.35 | 1.4 |
| R. Postcentral White Matter | 0.38 | 0.52 |
| R. Precentral White Matter | 0.38 | 0.60 |
| R. Supramarginal White Matter | 0.41 | 0.21 |
| R. Corona Radiata and Internal Capsule | 0.43 | 0.078 |

**Supplementary Table 2.**
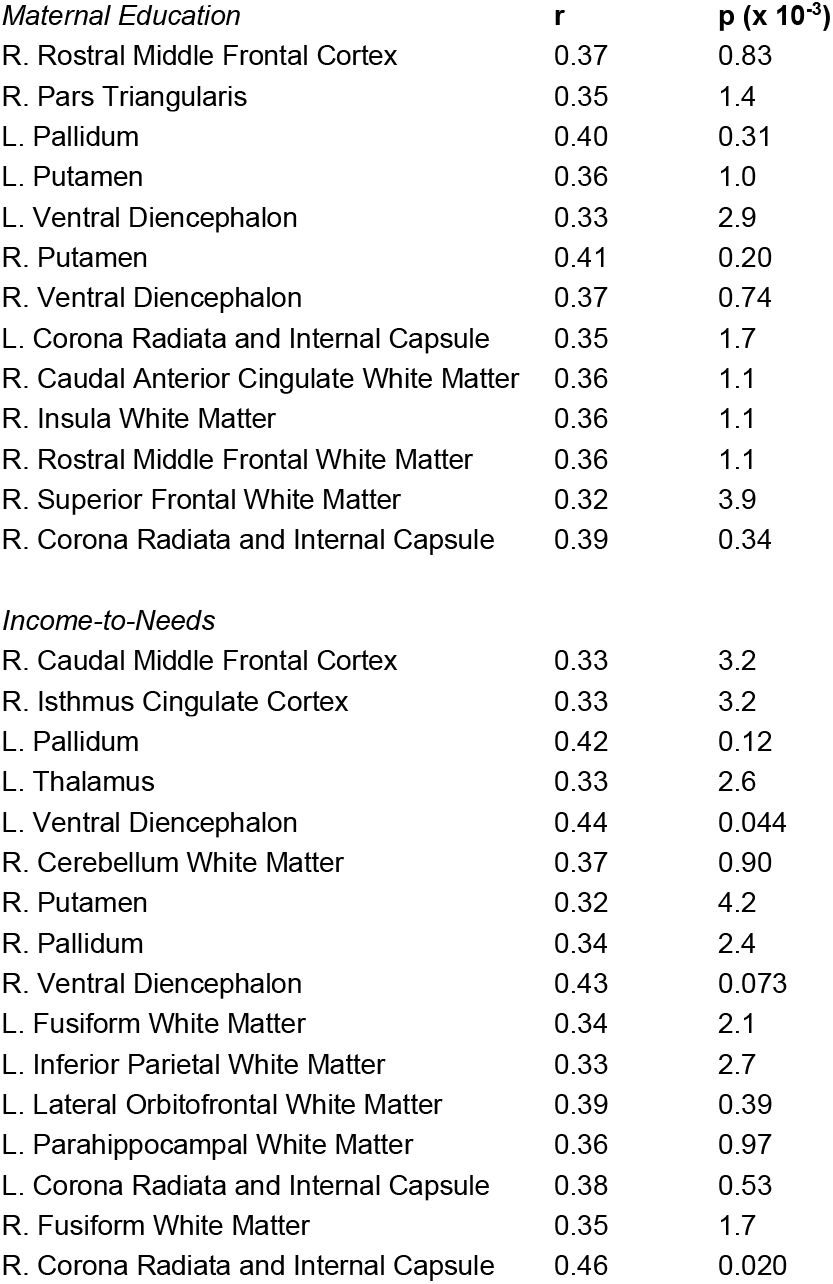
Brain-Poverty Relationships

**Supplementary Table 3.** Indirect Effects

| *Maternal Education → HAZ → Brain Volume* | Estimate | Lower CI | Upper CI | p |
| --- | --- | --- | --- | --- |
| R. Superior Temporal Cortex | 0.28 | 0.027 | 0.95 | 0.027 |
| L. Amygdala | 0.30 | 0.053 | 0.90 | 0.020 |
| L. Cerebellum White Matter | 0.37 | 0.067 | 1.5 | 0.017 |
| L. Pallidum | 0.28 | 0.065 | 0.92 | 0.0098 |
| R. Amygdala | 0.38 | 0.11 | 1.0 | 0.0092 |
| R. Cerebellum White Matter | 0.21 | 0.0067 | 0.72 | 0.041 |
| R. Pallidum | 0.38 | 0.067 | 1.5 | 0.021 |
| Corpus Callosum | 0.28 | 0.015 | 1.6 | 0.039 |
| L. Corona Radiata and Internal Capsule | 0.25 | 0.031 | 0.80 | 0.022 |
| R. Corona Radiata and Internal Capsule | 0.28 | 0.046 | 0.82 | 0.011 |
| *Maternal Education → WAZ → Brain Volume* |  |  |  |  |
| L. Cerebellum White Matter | 0.36 | 0.060 | 1.5 | 0.020 |
| L. Pallidum | 0.26 | 0.043 | 0.88 | 0.017 |
| R. Amygdala | 0.28 | 0.033 | 0.98 | 0.028 |
| R. Pallidum | 0.31 | 0.016 | 1.4 | 0.038 |
| Corpus Callosum | 0.28 | 0.022 | 1.6 | 0.035 |
| L. Corona Radiata and Internal Capsule | 0.27 | 0.045 | 0.89 | 0.015 |
| R. Corona Radiata and Internal Capsule | 0.28 | 0.045 | 0.85 | 0.013 |
| *Income-to-Needs → HAZ → Brain Volume* |  |  |  |  |
| L. Banks, Superior Temporal Sulcus | 0.31 | 0.053 | 1.2 | 0.021 |
| R. Isthmus Cingulate Cortex | 0.28 | 0.076 | 0.70 | 0.0044 |
| R. Superior Temporal Cortex | 0.33 | 0.070 | 0.98 | 0.0080 |
| L. Thalamus | 0.17 | -0.013 | 0.57 | 0.074 |
| R. Amygdala | 0.26 | 0.015 | 1.5 | 0.040 |
| Corpus Callosum | 0.32 | 0.11 | 0.76 | 0.0046 |
| L. Insula White Matter | 0.37 | 0.036 | 1.5 | 0.028 |
| L. Precentral White Matter | 0.24 | 0.045 | 0.64 | 0.012 |
| R. Isthmus Cingulate White Matter | 0.32 | 0.0091 | 1.2 | 0.043 |
| R. Precentral White Matter | 0.32 | 0.045 | 0.96 | 0.019 |
| R. Corona Radiata and Internal Capsule | 0.24 | 0.054 | 0.55 | 0.0054 |
| *Income-to-Needs → WAZ → Brain Volume* |  |  |  |  |
| L. Posterior Cingulate Cortex | 0.29 | 0.068 | 0.81 | 0.0054 |
| L. Cerebellum White Matter | 0.32 | 0.036 | 1.0 | 0.020 |
| R. Amygdala | 0.30 | 0.034 | 1.6 | 0.031 |
| L. Fusiform White Matter | 0.28 | 0.077 | 0.71 | 0.0052 |
| L. Precentral White Matter | 0.38 | 0.023 | 1.6 | 0.036 |
| L. Corona Radiata and Internal Capsule | 0.31 | 0.074 | 0.74 | 0.0066 |
| R. Corona Radiata and Internal Capsule | 0.29 | 0.066 | 0.64 | 0.0082 |
| *Income-to-Needs → WHZ → Brain Volume* |  |  |  |  |
|  | Estimate | Lower CI | Upper CI | p |
| L. Corona Radiata and Internal Capsule | 0.23 | 0.021 | 0.62 | 0.026 |
| R. Corona Radiata and Internal Capsule | 0.21 | 0.0071 | 0.52 | 0.041 |

## References

1. C. A. Nelson, L. J. Gabard-durnam, Early Adversity and Critical Periods: Neurodevelopmental Consequences of Violating the Expectable Environment. Trends Neurosci. 43, 133–143 (2020).

2. S. Grantham-McGregor, et al., Child development in developing countries. Lancet 369, 60–70 (2007).

3. C. C. John, M. M. Black, C. A. Nelson, Neurodevelopment: The Impact of Nutrition and Inflammation During Early to Middle Childhood in Low-Resource Settings. Pediatrics 139, S59–S71 (2017).

4. R. E. Black, et al., Maternal and child undernutrition and overweight in low-income and middle-income countries. Lancet 382, 427–451 (2013).

5. M. De Onis, F. Branca, Review Article Childhood stunting: a global perspective. Matern. Child Nutr. 12, 12–26 (2016).

6. A. E. Schnee, et al., Identification of Etiology-Specific Diarrhea Associated with Linear Growth Faltering in Bangladeshi Infants. Am. J. Epidemiol. 187, 2210–2218 (2018).

7. C. P. Stewart, L. Iannotti, K. G. Dewey, K. F. Michaelsen, A. W. Onyango, Contextualising complementary feeding in a broader framework for stunting prevention. Matern. Child Nutr. 9, 27–45 (2013).

8. Z. A. Bhutta, et al., Severe childhood malnutrition. Nat. Rev. Dis. Prim. 3, 1–18 (2017).

9. A. Fuglestad, R. Rao, M. Georgieff, M. Code, “The role of nutrition in cognitive development” in Handbook of Developmental Cognitive Neuroscience, C. Nelson, M. Luciana, Eds. (MIT Press, 2008), pp. 623–42.

10. W. Xie, et al., NeuroImage Chronic in fl ammation is associated with neural responses to faces in bangladeshi children. Neuroimage 202, 116110 (2019).

11. W. Xie, et al., Growth faltering is associated with altered brain functional connectivity and cognitive outcomes in urban Bangladeshi children exposed to early adversity. BMC Med. 17, 1–11 (2019).

12. C. Hayashi, et al., “Levels and trends in child malnutrition” (2018).

13. M. de Onis, A. Onyango, E. Borghi, A. Siyam, Worldwide implementation of the WHO Child Growth Standards. Public Health Nutr. 15, 1603–1610 (2012).

14. J. R. Donowitz, et al., Role of maternal health and infant inflammation in nutritional and neurodevelopmental outcomes of two-year-old Bangladeshi children. PLoS Negl. Trop. Dis. 12, 1–20 (2018).

15. S. Jensen, A. Berens, C. Nelson, Effects of poverty on interacting biological systems underlying child development. Lancet Child Adolesc. Heal. 1, 225–239 (2017).

16. T. Turesky, et al., Relating anthropometric indicators to brain structure in 2-month-old Bangladeshi infants growing up in poverty: A pilot study. Neuroimage 210, 1–10 (2020).

17. G. D. Gunston, D. Burkimsher, H. Malan, A. A. Sive, Reversible cerebral shrinkage in kwashiorkor: an MRI study. Arch. Dis. Child. 67, 1030–1032 (1992).

18. O. M. Atalabi, I. A. Lagunju, O. O. Tongo, O. O. Akinyinka, Cranial magnetic resonance imaging findings in Kwashiorkor. Int. J. Neurosci. 120, 23–27 (2010).

19. A. M. El-sherif, G. M. Babrs, A. M. Ismail, Cranial Magnetic Resonance Imaging (MRI) Changes in Severely Malnourished Children before and after Treatment. Life Sci. J. 9, 738–742 (2012).

20. H. Hulshoff, et al., Prenatal Exposure to Famine and Brain Morphology in Schizophrenia. Am. J. Psychiatry 157, 1170–1172 (2000).

21. M. J. H. Rytter, L. Kolte, A. Briend, H. Friis, V. B. Christensen, The immune system in children with malnutrition - A systematic review. PLoS One 9 (2014).

22. A. Jefferson, et al., Inflammatory biomarkers are associated with total brain volume The Framingham Heart Study. Neurology 68, 1032–1039 (2007).

23. P. J. Duggan, et al., Intrauterine T-cell activation and increased proinflammatory cytokine concentrations in preterm infants with cerebral lesions. Lancet 358, 1699–1700 (2001).

24. I. Hansen-Pupp, et al., Circulating interferon-gamma and white matter brain damage in preterm infants. Pediatr. Res. 58, 946–952 (2005).

25. B. H. Yoon, et al., Amniotic fluid inflammatory cytokines (interleukin-6, interleukin-1(beta), and tumor necrosis factor-(alpha)), neonatal brain white matter lesions, and cerebral palsy. Am. J. Obstet. Gynecol. 177, 19–26 (1997).

26. C. L. McDermott, et al., Longitudinally mapping childhood socioeconomic status associations with cortical and subcortical morphology. J. Neurosci. 39, 1365–1373 (2019).

27. K. G. Noble, S. M. Houston, E. Kan, E. R. Sowell, Neural correlates of socioeconomic status in the developing human brain. Dev. Sci. 15, 516–527 (2012).

28. K. G. Noble, et al., Family income, parental education and brain structure in children and adolescents. Nat. Neurosci. 18, 773–778 (2015).

29. J. Luby, et al., The effects of poverty on childhood brain development: The mediating effect of caregiving and stressful life events. JAMA Pediatr. 167, 1135–1142 (2013).

30. J. L. Hanson, A. Chandra, B. L. Wolfe, S. D. Pollak, Association between income and the hippocampus. PLoS One 6, 1–8 (2011).

31. M. J. Farah, The Neuroscience of Socioeconomic Status: Correlates, Causes, and Consequences. Neuron 96, 56–71 (2017).

32. P. Kim, C. Capistrano, C. Congleton, Socioeconomic disadvantages and neural sensitivity to infant cry: Role of maternal distress. Soc. Cogn. Affect. Neurosci. 11, 1597–1607 (2016).

33. P. Kim, et al., Effects of childhood poverty and chronic stress on emotion regulatory brain function in adulthood. Proc. Natl. Acad. Sci. 110, 18442–18447 (2013).

34. A. Klein, et al., Mindboggling morphometry of human brains (2017).

35. K. J. Gorgolewski, et al., BIDS apps: Improving ease of use, accessibility, and reproducibility of neuroimaging data analysis methods. PLoS Comput. Biol. 13, 1–16 (2017).

36. G. Lohmann, D. Y. Von Cramon, C. F. Colchester, Deep Sulcal Landmarks Provide an Organizing Framework for Human Cortical Folding. Cereb. Cortex 18, 1415–1420 (2008).

37. C. Victora, M. de Onis, P. Hallal, M. Blossner, R. Shrimpton, Worldwide Timing of Growth Faltering: Revisiting Implications for Interventions. Pediatrics 125, e473–e480 (2010).

38. S. Jensen, F. Tofail, R. Haque, W. Petri, C. Nelson, Child development in the context of biological and psychosocial hazards among poor families in Bangladesh. PLoS One 14, 1–17 (2019).

39. S. Jensen, et al., Neural correlates of early adversity among Bangladeshi infants. Sci. Rep. 9, 1–10 (2019).

40. G. Moreau, et al., Childhood Growth and Neurocognition are Associated with Distinct Sets of Metabolites Article. EBioMedicine in press (2019).

41. T. Turesky, et al., The relationship between biological and psychosocial risk factors and resting-state functional connectivity in 2-month-old Bangladeshi infants: a feasibility and pilot study. Dev. Sci. 22, e12841 (2019).

42. M. de Onis, C. Garza, C. G. Victora, M. K. Bhan, K. R. Norum, The WHO Multicentre Growth Reference Study (MGRS): Rationale, planning, and implementation. Food Nutr. Bull. 25, S3–S89 (2004).

43. M. de Onis, C. Garza, A. Onyango, R. Martorell, WHO child growth standards. Acta Paediatr. 95, 5–6 (2006).

44. J. D. Hamadani, et al., Critical windows of exposure for arsenic-associated impairment of cognitive function in pre-school girls and boys: A population-based cohort study. Int. J. Epidemiol. 40, 1593–1604 (2011).

45. M. Kippler, et al., Early-life cadmium exposure and child development in 5-year-old girls and boys: a cohort study in rural Bangladesh. Child. Heal. 120, 1462–1468 (2012).

46. A. J. Sameroff, R. Seifer, A. Baldwin, C. Baldwin, Stability of Intelligence from Preschool to Adolescence: The Influence of Social and Family Risk Factors. Soc. Res. child Dev. 64, 80–97 (1993).

47. N. M. Raschle, et al., Making MR imaging child’s play - Pediatric neuroimaging protocol, guidelines and procedure. J. Vis. Exp. 29, 1–6 (2009).

48. O. Esteban, et al., fMRIPrep: a robust preprocessing pipeline for functional MRI. Nat. Methods 16, 111–116 (2019).

49. Y. Benjamini, Y. Hochberg, Controlling the false discovery rate: a practical and powerful approach to multiple testing. J.R. Stat. 57, 289–300 (1995).

50. A. F. Hayes, Beyond Baron and Kenny: Statistical mediation analysis in the new millennium. Commun. Monogr. 76, 408–420 (2009).

51. G. Favrais, et al., Systemic Inflammation Disrupts the Developmental Program of White Matter. Ann. Neurol. 70, 550–65 (2011).

52. N. Makris, et al., Decreased Volume of the Brain Reward System in Alcoholism. Biol. Psychiatry 64, 192–202 (2008).

53. K. Keunen, R. M. Van Elburg, F. Van Bel, M. J. N. L. Benders, Impact of nutrition on brain development and its neuroprotective implications following preterm birth. Pediatr. Res. 77, 148–155 (2015).

54. G. De Palma, S. M. Collins, P. Bercik, E. F. Verdu, The microbiota-gut-brain axis in gastrointestinal disorders: Stressed bugs, stressed brain or both? J. Physiol. 592, 2989–2997 (2014).

55. M. R. Irwin, S. W. Cole, Reciprocal regulation of the neural and innate immune systems. Nat. Rev. Immunol. 11, 625–632 (2011).

56. N. Sudo, “Microbiome, HPA axis and production of endocrine hormones in the gut” in Microbial Endocrinology: The Microbiota Gut-Brain Axis in Health and Disease, M. Lyte, J. Cryan, Eds. (Springer, 2014), pp. 177–194.

57. S. J. Lupien, B. S. McEwen, M. R. Gunnar, C. Heim, Effects of stress throughout the lifespan on the brain, behaviour and cognition. Nat. Rev. Neurosci. 10, 434–445 (2009).

58. F. A. Middleton, P. L. Strick, Anatomical evidence for cerebellar and basal ganglia involvement in higher cognitive function. Science (80-.). 266, 458–461 (1994).

59. N. L. Hair, J. L. Hanson, B. L. Wolfe, S. D. Pollak, Association of child poverty, brain development, and academic achievement. JAMA Pediatr. 169, 822–829 (2015).

60. R. K. Lenroot, et al., Sexual dimorphism of brain developmental trajectories during childhood and adolescence. Neuroimage 36, 1065–1073 (2007).

61. C. E. Sanchez, J. E. Richards, C. R. Almli, Neurodevelopmental MRI brain templates for children from 2 weeks to 4 years of age. Dev. Psychobiol. 54, 77–91 (2012).

62. J. Giedd, et al., Brain development during childhood and adolescence: a longitudinal MRI study. Appl. Phys. Lett. 2, 861–863 (1999).

63. K. Jednorog, et al., The Influence of Socioeconomic Status on Children ‘ s Brain Structure. PLoS One 7, 1–9 (2012).

64. N. H. Brito, K. G. Noble, Socioeconomic status and structural brain development. Front. Neurosci. 8, 1–12 (2014).

65. C. A. Nelson, Hazards to Early Development: The Biological Embedding of Early Life Adversity. Neuron 96, 262–266 (2017).

66. Cell Editorial Team, Science Has a Racism Problem. Cell 181, 1443–1444 (2020).

